# Optogenetic activation of nonhuman primate cortical and subcortical brain circuits highlights detection capabilities of MEG source imaging

**DOI:** 10.1101/2020.08.20.259465

**Authors:** GE Alberto, JR Stapleton-Kotloski, DC Klorig, ER Rogers, C Constantinidis, JB Daunais, DW Godwin

**Affiliations:** Wake Forest School of Medicine Department of Neurobiology and Anatomy; Wake Forest School of Medicine Department of Neurology; Wake Forest School of Medicine Department of Physiology and Pharmacology; Research and Education Department, W.G. (Bill) Hefner Veterans Affairs Medical Center

## Abstract

Magnetoencephalography (MEG) measures neuromagnetic activity with high temporal, and theoretically, high spatial resolution. However, the ability of magnetic source imaging (MSI) to localize deep sources is uncertain. We developed an experimental platform combining MEG-compatible optogenetic techniques in non-human primates (NHPs) to test the ability of MEG/MSI to image deep signals. We demonstrate localization of optogenetically-evoked signals to known sources in the superficial arcuate sulcus of cortex and in CA3 of hippocampus at a resolution of 750 µm^3^. In response to stimulation of arcuate sulcus and hippocampus, we detected activation in subcortical and thalamic structures, or extended temporal networks, respectively. This is the first demonstration of accurate localization of deep sources within an intact brain using a novel combination of optogenetics with MEG/MSI. This approach is suitable for exploring causal relationships between discrete brain regions through precise optogenetic control and simultaneous whole brain recording.

## INTRODUCTION

A fundamental goal of neuroimaging is to observe and quantify brain activity across the spatial and temporal resolutions at which it occurs. MEG holds particular promise for improving upon current neuroimaging methods because it has identical temporal resolution to EEG, with the potential for superior spatial resolution^1,2^. MEG detects biomagnetic fluctuations that co-occur with neuronal activity. Unlike electric fields, biomagnetic signals are not distorted by the intervening tissues of the head prior to arriving at the sensor thereby providing the estimation of the source of these signals with magnetic source imaging (MSI) an inherent advantage over source localization with EEG^2^. While MEG has been employed clinically for use in the pre-surgical evaluation of epilepsy^7–10^ and in studies of deep structures such as hippocampus^11^ and amygdala^12^, a detailed investigation of the resolution capabilities of MEG is currently lacking.

Early studies employing simultaneous intracranial EEG and MEG in patients with epilepsy suggested that MEG was unable to detect activity originating in deep structures such as the hippocampus^13–15^. Possible explanations for this failure included concerns that the curved architecture of the hippocampus caused magnetic fields to cancel or that the distance from the MEG sensors to hippocampus may be too great^16^. Theoretically however, the combination of a larger number of axial gradiometers (shown to have a superior depth profile in comparison to planar gradiometers^4,16^), a low noise floor, minimal motion^5,6^, and appropriate use of beamformers^3,8,11^ should optimize the signal-to-noise ratio (SNR) of biomagnetic signals to allow sub-mm^3^ localization of activity^5^ throughout the volume of the brain. However, without unequivocal demonstration in an *in vivo* preparation the degree to which the theoretical capabilities of MSI are reflected in practice remains uncertain.

We have developed a MEG-compatible optogenetic preparation in vervet monkeys that allows simultaneous optical stimulation and MEG recordings, along with associated local field potential (LFP) recordings from deep and superficial structures. Stimulation and LFP recordings were achieved with an optrode, which is a combined optical fiber/recording electrode^20, 50^. Optogenetic techniques are well suited for use with MEG/MSI in that they are magnetically silent and allow for precise control over neural population potentials through the use and activation of virally-expressed, light sensitive ion channels^17^. In this report we have relied on synthetic aperture magnetometry (SAM)^3^, a linearly-constrained minimum variance (LCMV) beamformer method^3,8,11^ with a theoretical sub-mm^3^ spatial resolution^4–6^. We have used SAM to convert MEG signals into whole-brain statistical parametric maps (SPMs), differentiating it from more traditional EEG and MEG dipole analyses^3^. Using this combination of approaches, we elicited and measured both biomagnetic and electrophysiological activity in two representative brain areas, the arcuate sulcus of the cerebral cortex and within the CA3 region of the hippocampus in vervet monkeys. We demonstrate the ability to selectively stimulate and detect neuronal activity in these areas, as well as to accurately localize discrete optogenetically-elicited sources of activity in brain regions downstream from stimulated areas. We further demonstrate the utility of combining optogenetics and MEG/MSI in experimenter-controlled functional brain mapping across a wide range of spatiotemporal scales. We propose that the tight temporal and spatial control over neural populations afforded with optogenetics coupled with whole brain, high spatiotemporal resolution MEG recordings will enable a new approach to functional neuroimaging.

## RESULTS

### Electrophysiology of Early Expression

Three vervet monkeys were injected with AAV10-CaMKIIa-ChR2-eYFP and an optrode was implanted at the site of transfection. After the animals had recovered, it was determined whether there was a detectable LFP via the indwelling optrode during the early expression period. All reported experimentation and data acquisition occurred under anesthesia. Optical pulses were delivered through the implanted fiber at both the cortical and hippocampal sites to test the response of newly transfected tissue to stimulation. Functional expression was determined using optogenetic intensity-response curves at each site 2, 5, and 7 weeks post-injection. An optogenetically evoked potential was detectable from depth recordings by 5 weeks post-expression and had stabilized by ∼7 weeks (Figure 1c-e; cortical: baseline (2 week) 95% confidence interval (CI) [-0.151, 0.374], post-expression (5 week) 95% CI [0.522, 1.12], post-expression (7 week) 95% CI [0.683, 1.09], hippocampal: baseline (2 week) 95% CI [-4.55e-3, 0.455], post-expression (5 week) 95% CI [1.31, 1.65], post-expression (7 week) 95% CI [1.50, 1.89]).

**Figure 1.**
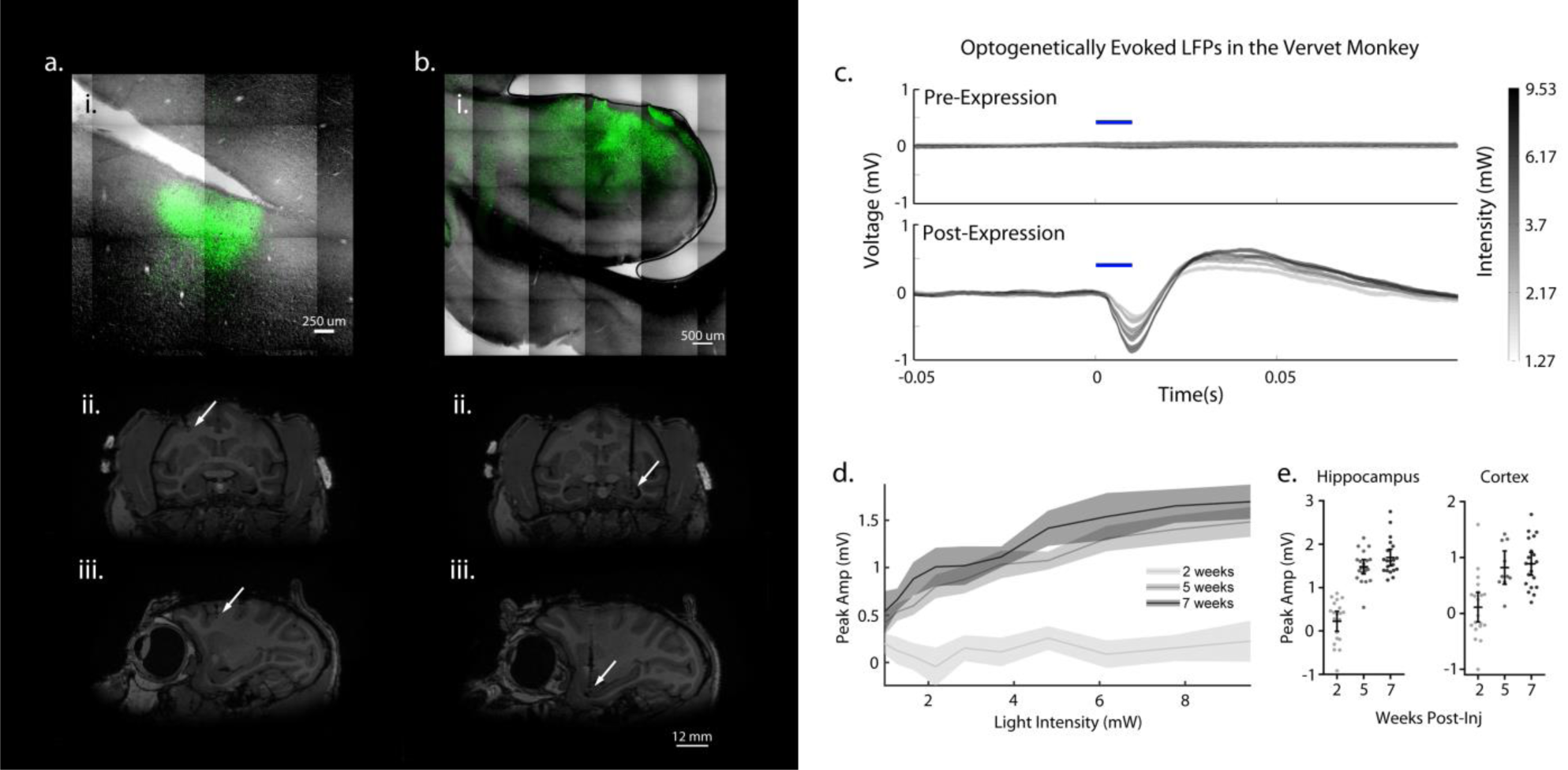
Location of optogenetic protein expression and light evoked responses. **ai**. ChR2-eYFP labeled neurons confined to gray matter of the posterior wall of right arcuate sulcus corresponding to area 8a, white line indicates approximate boundary between gray and white matter. Coronal (**ii**) and sagittal (**iii**) MRI demonstrating colocalization of the optrode. **bi**. ChR2-eYFP labeled neurons in the anterior dorsolateral aspect of left hippocampus corresponding to area CA3 indicate transgene expression. Coronal (**ii**) and sagittal (**iii**) MRI demonstrating colocalization of the optrode. **c**. Example of averaged response to stimulation at five intensity levels (grey to black) in hippocampus showing absence of response in week 2 (upper trace) compared to the large, biphasic LFP in week 7 (lower trace). Blue bar indicates stimulus onset and duration. **d**. Average intensity-response curves for each of the three time points. Bands = 95% CI. **e**. Maximum intensity stimulation (9.53 mW) of cortex and hippocampus at 7 weeks post transfection elicited a response as measured by the LFP electrode that was greater than at 2 weeks post-transfection. AS= arcuate sulcus; GM= gray matter; WM = white matter; DG= dentate gyrus; SB= subiculum.

### MEG Recordings of Early Expression

We analyzed post-surgical (2 week) MEG data using SAM. Consistent with the electrophysiological recordings obtained in the same week, there were no detectable changes (local maxima or minima) in the SAM statistical parametric maps (SPMs) at the site of stimulation, and the maps were qualitatively similar to the pre-surgical control scans (Not shown - see Methods for beamforming parameters). We conclude that expression of ChR2 at two weeks post-injection was insufficient to generate an optically evoked response that could be identified using SAM, further evidence that our preparation is insensitive to light stimulation in the absence of ChR2.

Post-mortem, fluorescence confocal microscopy revealed ChR2-eYFP at the targeted sites of transfection, indicating uptake of the vector by the surrounding neurons and expression of the transgene. Labeled neurons were observed in the posterior wall of arcuate cortex corresponding to area 8a of the monkey cortex^23,24^ (Figure 1ai). Hippocampal imaging revealed ChR2-eYFP labeled neurons in anterior dorsal hippocampus corresponding to area CA3^24^ with labeled efferent connections observable in the surrounding tissue (Figure 1bi). Structural MRI prior to necropsy confirmed placement of the optrode in both the posterior wall of arcuate sulcus and in anterior dorsal hippocampus^25^ (Figures 1aii-iii and Figures 1bii-iii).

As a positive control for MSI detection and localization accuracy, we presented 119, 50 ms pulses of white light to the left lower quadrant of the visual field of the left eye. (The right eye was closed.) In contrast to the lack of optogenetic biomagnetic responses at 2 weeks, visual stimulation experiments conducted at this time point demonstrated SAM peak voxels in the right occipital lobe (±50 ms map window relative to stimulus onset, bandwidth of 20-35 Hz, pseudo-t = 0.3) consistent with the early cortical component of visual evoked potentials^22^ (Figure S1aviii). In addition to activation in occipital cortex, SAM also revealed peaks associated with the visual network, including (aiii.) left optic tract (±75 ms map, 12-35 Hz bandwidth, pseudo-t = 0.4), (aiv.) left lateral geniculate n. (±50 ms map, 20-50 Hz bandwidth, pseudo-t = 0.4), and (avii.) left superior colliculus (±50 ms map, 20-60 Hz, pseudo-t = 0.4. Additional peaks were seen in (aii.) left anterior commissure (±75 ms map, 12-35 Hz bandwidth, pseudo-t = 0.4), (av.) right posterior hippocampus (±75 ms map, 12-55 Hz bandwidth, both pseudo-t’s = 0.3), (avi.) right posterior white matter tracts (±75 ms map, 12-60 Hz bandwidth, pseudo-t = 0.4), and (ai.) right arcuate cortex (±75 ms map, 12-120 Hz bandwidth, pseudo-t = 0.6.). Importantly, right arcuate and left hippocampal responses were also detected in the LFP in response to visual stimulation (Figure S1b). The visual stimulation experiment, in addition to serving as a positive control for evoked signals, also demonstrates that SAM can localize primary sensory cortex in response to stimulation, that it can reveal subcortical brain structures known to be involved in the processing of visual stimulation, and that activated white matter tracts may also localized with SAM.

### MEG Recordings of Stable Expression

Eight weeks after transfection, and once the neural response to optical stimulation had reached a plateau, MEG recordings commenced. Figure 2a depicts an example of optogenetic activation (50 ms square pulses) in one NHP as localized by dual-state SAM to the posterior bank of the right arcuate sulcus in the coronal (i), sagittal (ii), and axial planes (iii), as well as additional cortical peaks (local maxima or minima in the SAM SPMs, Figure 2aii) near the site of optical stimulation. Peaks in the SAM SPMs (local maxima or minima in the maps, here in units of pseudo-t scores) can be interpreted as significant voxels in comparison to baseline and represent voxels of highly synchronous (red) or desynchronous (blue) activity within the frequency band of interest. All brain images are depicted in radiological coordinates, in which the right hemisphere is presented on the left. The whole brain, un-thresholded, dual-state SAM SPM is depicted in Figure 2aiv; note the region of synchronized activity (red voxels) arising from the site of activation and surrounded by desynchronized activity (blue voxels). For Figures 2ai-iv, the dual-state beamforming parameters for the map were a ±75 ms window relative to stimulus onset, for a frequency band of 15-35 Hz, and resulted in a pseudo-t score of 0.6. Figure 2b depicts the SAM arcuate peaks corresponding to four additional and different optical stimulus inputs. Figures 2bi-ii present peak activations in right arcuate for one NHP (same subject as in Figure 2a) and Figures 2biii-iv shows right arcuate stimulation for a second NHP. Figure 2bi depicts the arcuate peak (mapped at a ±3.5 s window relative to stimulus onset, bandwidth of DC-20 Hz, pseudo-t = 0.5) in response to 8 Hz, sine waves (i.e., sinusoidal-modulation of the LED), and Figure 2bii shows the arcuate peak in response to 40 Hz sine waves (±3.5 s map window, bandwidth of 20-50 Hz, pseudo-t = 0.7). The results for arcuate stimulation in a second NHP are presented in Figures 2biii-iv; Figure 2biii shows the SAM arcuate peak in response to 10 ms square wave pulses (±600 ms map, bandwidth of 3-70 Hz, pseudo-t = 0.2), whereas Figure 2biv exhibits arcuate activity elicited by 20 Hz square wave pulse trains (5 ms width for each square wave; ±200 ms map, bandwidth of 5-70 Hz, pseudo-t = 0.3). Details on the stimuli and beamformer parameters for each stimulus type are presented in Methods. MSI can also be used to construct virtual electrodes (source series) for any voxel in the brain, providing a continuous, wide-band, sub-millisecond readout of activity that may provide similar neurophysiological data relative to that resulting from an actual invasive electrode^7^. Figure 2bv presents a single trial of a local field potential (white) recorded from the indwelling optrode and a simultaneous SAM virtual electrode (red, viewing filters set from 1-30 Hz) extracted from the arcuate peak shown in Figure 2biii. The blue vertical line indicates the time of stimulus onset, in this case a 10 ms square wave pulse. Both the LFP and virtual electrode exhibit a rapid rise and peak in activity following stimulus onset, and both qualitatively share similar features and time courses on a single trial basis.

**Figure 2.**
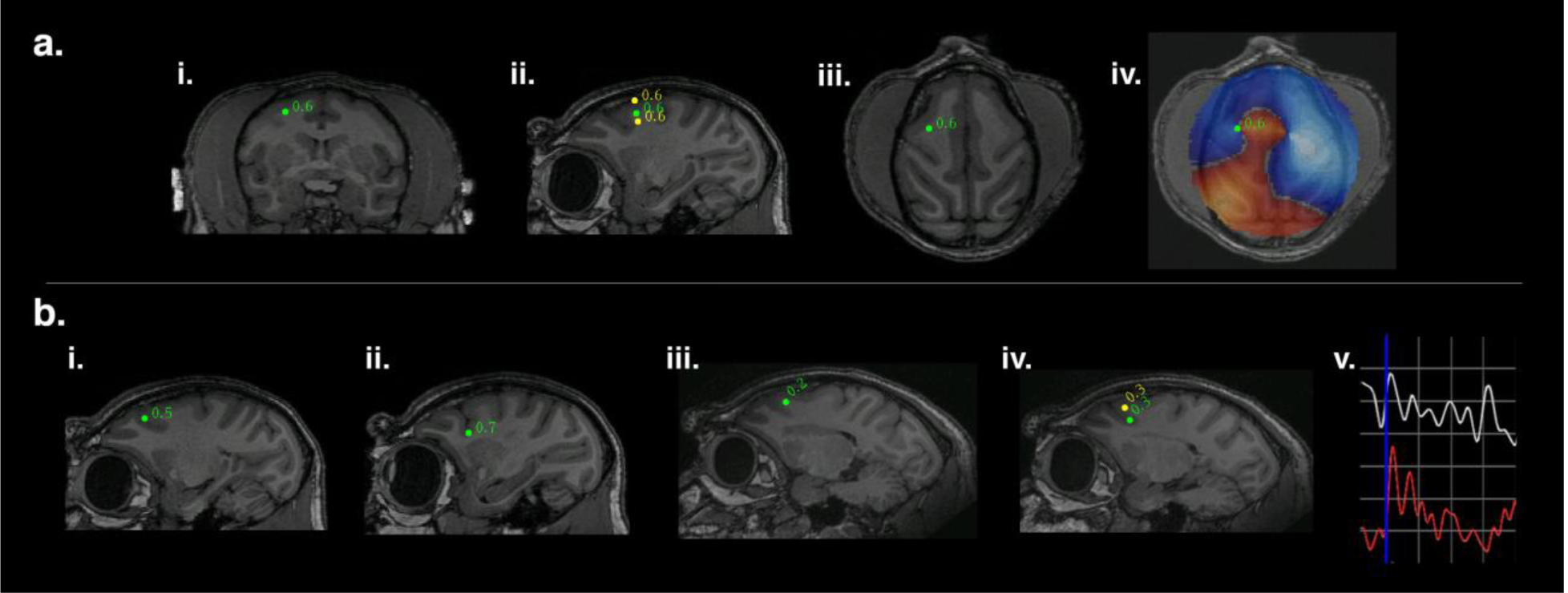
SAM source localization of optogenetically evoked MEG responses in the right arcuate sulcus. **a**. An example of optogenetic stimulation localized by dual-state SAM to the posterior bank of right arcuate sulcus in the coronal (**i**), sagittal (**ii**), and axial planes (**iii**), for a stimulus of 50 ms square light pulses. The whole brain, un-thresholded, dual-state SAM SPM (voxel size of 750 µm^3^, **iv**) reveals synchronized activity (red voxels) arising from the stimulation site and surrounded by desynchronization (blue voxels). Green dot and number indicate a peak in the SPM plus the associated pseudo-t value. **b**. Right SAM arcuate peaks for four additional and different optical stimuli in two different NHPs. **bi-ii** present arcuate activations for one NHP and **iii-iv** shows arcuate stimulation for a second NHP. **bi-ii** is the same subject as presented in **ai-iv**; **i** depicts the arcuate peak for 8 Hz sine waves, and **ii** shows the arcuate peak for 40 Hz sine waves. Arcuate stimulations in a second NHP are presented in **iii-iv**; **iii** shows the arcuate peak for 10 ms square pulses and **iv** shows the arcuate peak for 20 Hz square wave pulse trains. A single trial (**v**) of an LFP recorded by the optrode and a simultaneous SAM virtual electrode for an arcuate peak (as seen in **biii**). Blue vertical line indicates stimulus onset (10 ms pulses). Both the LFP (white) and virtual electrode (red) exhibit a rapid peak following stimulation and have similar features and time courses. One gray square in the graph = 100 ms on the abscissa and for the ordinate, 35 μV for the LFP or 15 nA-m for the virtual electrode. All maps follow radiological conventions.

Consistent activity was also elicited in the left hippocampi of two NHPs in response to a variety of optical stimulus types. Figure 3a presents an example of the left SAM hippocampal peak co-localized to the site of optical stimulation in a third subject, in this case a single, 60 s duration, 20 Hz sine wave. Figures 3ai-iii depict the peak in the coronal (i), sagittal (ii), and axial planes (iii), and Figure 3aiv illustrates the un-thresholded, whole-brain SAM dual state map (±4 s window, 10-30 Hz band, pseudo-t = 1.7) associated with the 20 Hz sine wave input. Note the narrow band of synchronization (red) confined to the dorsal aspect of the left hippocampus, and synchronous activity propagating towards left insula (e.g. see Figure 5a). The second row of Figure 3 depicts the left hippocampal peaks elicited by additional optogenetic stimulation for two different NHPs. The first two coronal slices (Figures 3bi-ii) correspond to hippocampal recordings obtained from a third NHP (same subject as presented in Figure 3a), and the last two coronal slices depict activity for the first NHP, the latter for whose arcuate peaks were depicted in Figure 2a. For Figure 3bi the stimulation parameters were single triangular (sawtooth) waves (±150 ms windows, 8-45 Hz band, pseudo-t = 0.8); for Figure 3bii the parameters were 10 ms square wave pulses (±150 ms windows, bandwidth of 7-85 Hz, pseudo-t = 0.2). Figure 3bv presents a single trial of a local field potential and the simultaneous SAM virtual electrode (viewing filters set from 1-35 Hz) extracted from the hippocampal peak shown in Figure 3bi. The blue vertical line indicates the time of the optical sawtooth onset. Both the LFP and virtual electrode exhibit a rapid rise and peak in activity following stimulus onset, and both qualitatively share similar features and time courses on a single trial basis. Hippocampal responses for the first NHP are depicted in Figure 3biii-iv with a hippocampal peak elicited by 8 Hz sine waves (±2 s windows, DC-15 Hz band, pseudo-t = 1.0, Figure 3biii), and another hippocampal peak elicited by 40 Hz sine waves (±2 s windows, 20-55 Hz band, pseudo-t = 0.6) shown in Figure 3biv.

**Figure 3.**
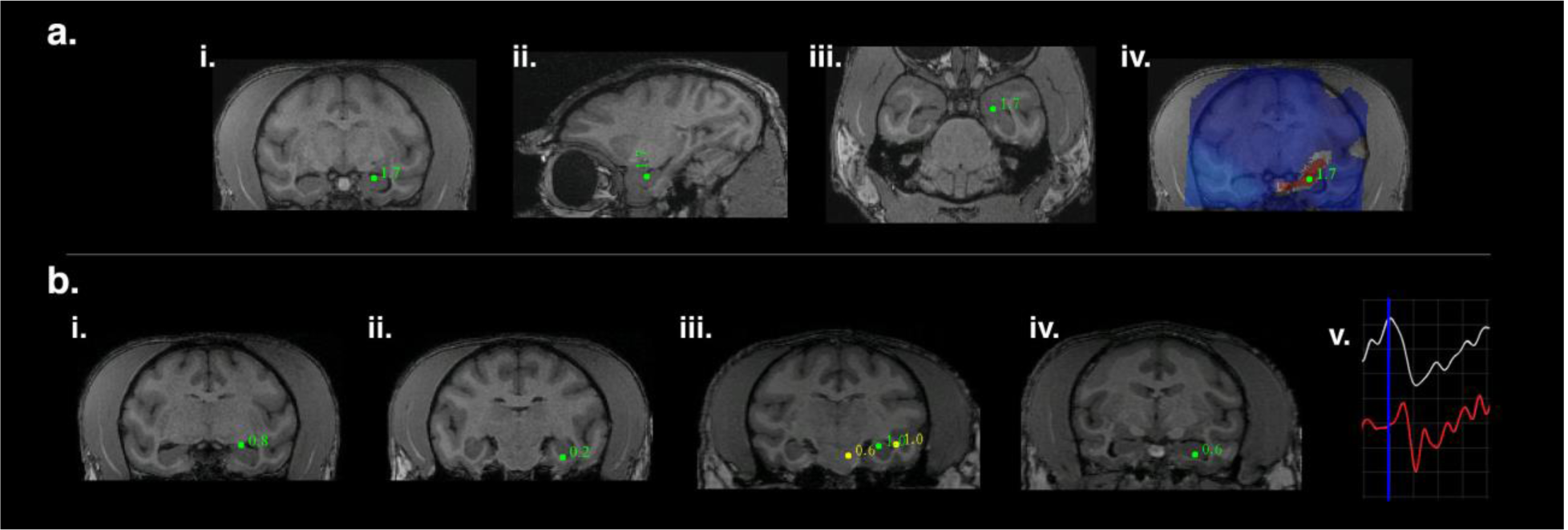
SAM source localization of optogenetically evoked MEG responses in the left hippocampus. **a**. SAM left hippocampal peak in coronal (**i**), sagittal (**ii**), and axial planes (**iii**) for a 20 Hz sine wave input in a third NHP. The whole brain, un-thresholded, SAM SPM shows synchronized activity arising from the stimulation site and surrounded by desynchronization (**iv**). **b**. Left SAM hippocampal peaks for four additional and different stimuli in two different NHPs. **bi-ii** is the same subject as presented in **a**; **i** and **ii** show hippocampal peaks for single 150 ms sawtooth trials or 10 ms square pulses, respectively. **biii-iv** show hippocampal peaks for 8 Hz or 40 Hz sine waves, respectively, in the same subject as in **Figure 2a**. A single trial (**v**) of a simultaneous LFP and SAM virtual electrode for hippocampus (as seen in **bi**). Blue vertical line indicates stimulus onset (sawtooth inputs). Both the LFP (white) and virtual electrode (red) peak rapidly following stimulation and have similar features and time courses. One gray square = 50 ms (abscissa) and 5 V for the LFP (amplified) or 500 nA-m for the virtual electrode (ordinate).

**Figure 4.**
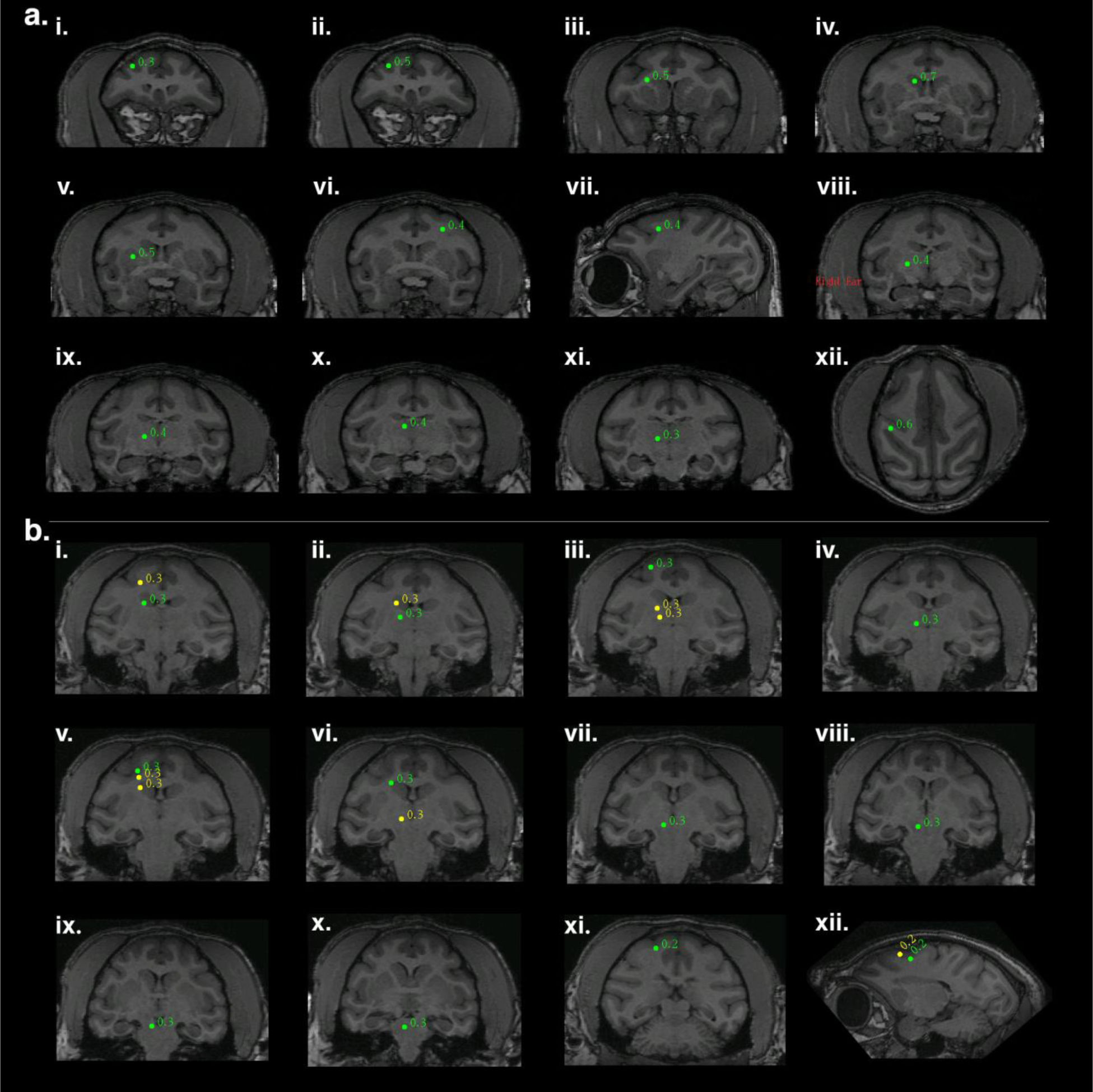
Arcuate optogenetic stimulation results in propagation through downstream networks. **a**. Identification of cortico-striato-pallido-thalamo-cortical (CSPTC) network activity in response to 8 Hz sine wave stimulation of right arcuate sulcus. SAM analysis of optical input to arcuate sulcus reveals a known functional network with peaks in: **i**. right inferior bank arcuate sulcus, **ii**. right superior bank arcuate sulcus, the site of stimulation, **iii**. right corona radiata/ dorsolateral caudate n., **iv**. right caudate n. **v**. right putamen/ external segment of globus pallidus, **vi**. and **vii**. posterior bank of contralateral (left) arcuate sulcus, coronal and sagittal slices, respectively, **viii**. right globus pallidus, **ix**. right ventroposterior medial / ventroposterior lateral n. of thalamus, **x**. right medial dorsal n. of thalamus, **xi**. right ventral posterior thalamus, and **xii**. parietal-occipital association area of the intraparietal sulcus. **b**. A second example of right hemispheric CSPTC network activity in a different NHP in response to 20 Hz square wave pulse trains with SAM peaks as follows: **i**. caudate n. (green) and cortical area 3 (yellow), **ii**. thalamus (green) and caudate n. (yellow), **iii**. posterior dorsal bank of the arcuate sulcus (green) and two thalamic peaks (yellow), **iv**. thalamus, **v**. posterior ventral bank of arcuate sulcus (green, site of stimulation), corpus callosum (yellow), and caudate n. (yellow), **vi**. caudate n. (green) and mesencephalic nuclei (yellow), **vii**. mesencephalic n., **viii**. substantia nigra, **ix**. tegmental n., **x**. pontine or mesencephalic n., and **xi**., and **xii**. all peaks in primary motor cortex (area 4).

**Figure 5.**
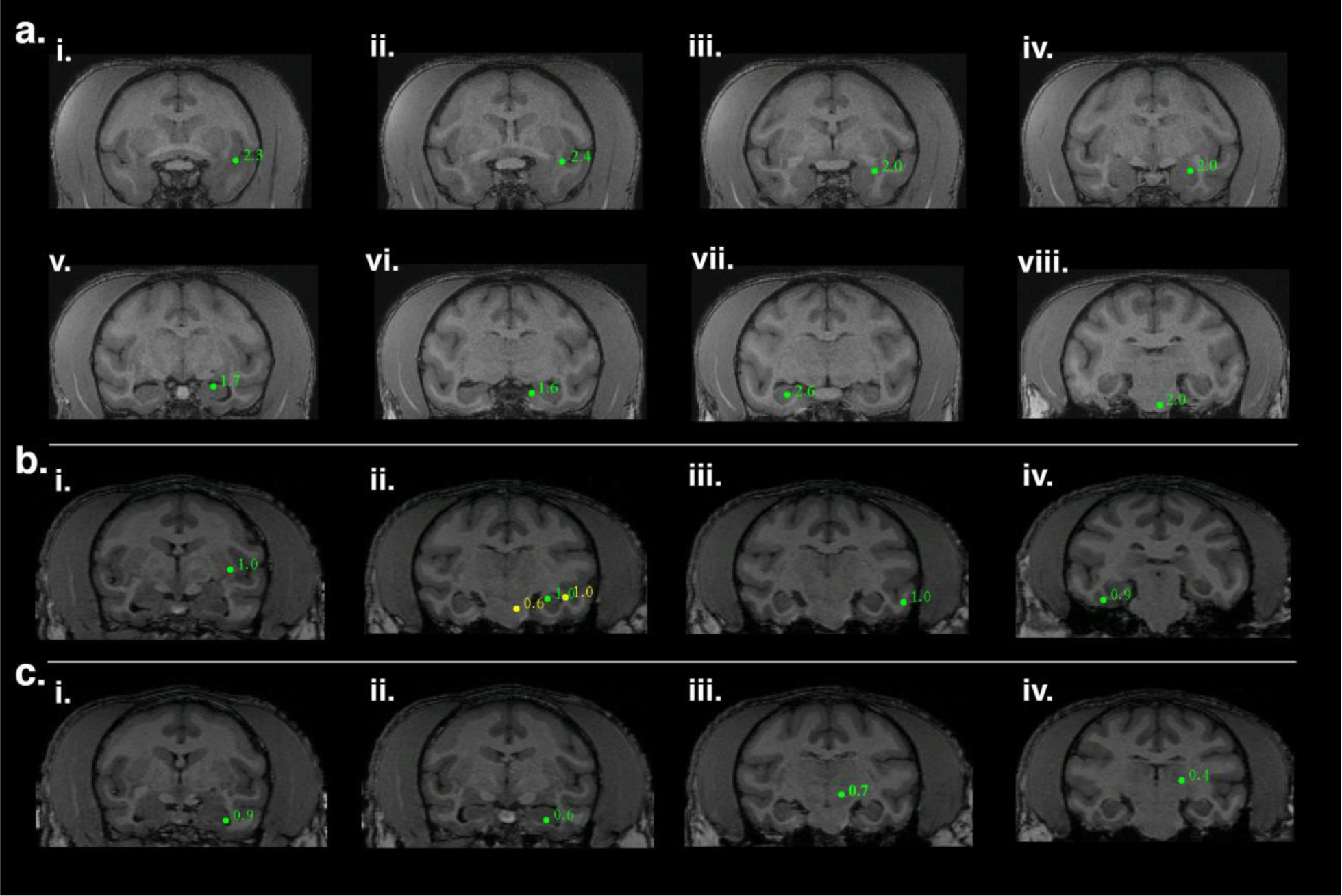
Hippocampal optogenetic stimulation results in propagation through downstream networks. **a**. Functional hippocampal and temporal network engaged by a 20 Hz optical sine input to left hippocampus (**v**) with SAM peaks as follows: **i**. left insular proisocortex/temporopolar proisocortex, **ii**. left para-insular cortex, **iii**. left dorsolateral amygdala (lateral n.), **iv**. left basolateral/lateral amygdala, **v**. left hippocampus, site of stimulation, **vi**. left parasubiculum/presubiculum of hippocampus, **vii**. contralateral (right) hippocampus, and **viii**. left pontine reticular activating formation. **b**. A second example of hippocampal network activity in a different NHP in response to 8 Hz sine wave input with SAM peaks as follows: **i**. left putamen, **ii**. PGa/IPa association areas of left temporal cortex (lateral yellow peak), left hippocampus (green peak, site of stimulation), and possible left red n. (yellow, medial peak), **iii**. area TEA/TEM of left temporal cortex, and **iv**. area TL/TFM of right temporal cortex. **c**. A third example of hippocampal network activity in response to 40 Hz sine wave input (same subject as in **b**.) with SAM peaks as follows: **i**. left anterior hippocampus, **ii**. left hippocampus, site of stimulation, **iii**. left deep mesencephalic n., and **iv**. left pulvinar n. of thalamus.

### Functional Mapping

In addition to eliciting biomagnetic activity at the transfection site, optogenetic stimulation also resulted in propagation through well-described downstream networks. Because the SAM maps were constructed for the whole brain volume, additional peaks were revealed in the various time-frequency SPMs. Interestingly, right arcuate stimulation elicited additional SAM peaks in a pattern that conformed to a known motor-associative network, the cortico-striatal-pallido-thalamocortical (CSPTC) motor network^26,27^. This network was revealed in both NHPs who received arcuate stimulation across different optical stimuli. An example of the CSPTC maps obtained for the first subject (as seen in Figure 2bi) are shown in Figure 4a as a function of 8 Hz sine wave input. In addition to the activation of right arcuate sulcus (ii), SAM maps also exhibited peaks in (i.) right anterior bank of the arcuate sulcus (±1 s windows, DC-12 Hz band, pseudo-t = 0.3), (iii.) right corona radiata/ dorsolateral tip of the caudate n. (±4 s windows, DC-15 Hz band, pseudo-t = 0.5), (iv.) right caudate n. (±2.5 s windows, DC-50 Hz band, pseudo-t =0.7), (v.) right putamen/ external segment of globus pallidus (±4 s windows, DC-15 Hz band, pseudo-t = 0.5), vi. and vii. posterior bank of contralateral (left) arcuate sulcus, coronal and sagittal slices, respectively (±3 s windows, DC-15 Hz band, pseudo-t = 0.4), (viii.) right globus pallidus (±4 s windows, DC-15 Hz band, pseudo-t = 0.4), (ix.) right ventroposterior medial / ventroposterior lateral n. of thalamus (±4 s windows, DC-15 Hz band, pseudo-t = 0.4), (x.) right medial dorsal n. of thalamus (±4 s windows, DC-15 Hz band, pseudo-t = 0.4), (xi.) ventral posterior thalamus (±3.5 s windows, DC-20 Hz band, pseudo-t = 0.3), and (xii.) parietal-occipital association area of the intraparietal sulcus (±2 s windows, DC-50 Hz band, pseudo-t = 0.6). A second example of right hemispheric CSPTC network activity for a different NHP in response to 20 Hz square wave pulse trains is presented in Figure 4b. Owing to the contrast of this subject’s MRI, the thalamic peak identities were less discernible. Despite this, SAM maps revealed peaks in (i.) caudate n. (green, ±200 ms windows, 5-70 Hz band, pseudo-t = 0.3) and corona radiata (yellow, ±200 ms windows, 5-70 Hz band, pseudo-t = 0.3), ii. thalamus (green, ±200 ms windows, 5-70 Hz band, pseudo-t = 0.3) and caudate n. (yellow, ±200 ms windows, 5-70 Hz band, pseudo-t = 0.3), iii. posterior dorsal bank of the arcuate sulcus (green, ±200 ms windows, 5-70 Hz band, pseudo-t = 0.3) and two thalamic peaks (yellow, both ±200 ms windows, 5-70 Hz band, pseudo-t’s = 0.3), iv. thalamus (±200 ms windows, 5-70 Hz band, pseudo-t = 0.3), v. posterior ventral bank of arcuate sulcus (green, site of stimulation, ±200 ms windows, 5-70 Hz band, pseudo-t = 0.3), corpus callosum (yellow, ±200 ms windows, 5-70 Hz band, pseudo-t = 0.3), and caudate n. (yellow, ±200 ms windows, 5-70 Hz band, pseudo-t = 0.3), vi. caudate n. (green, ±200 ms windows, 5-70 Hz band, pseudo-t = 0.3) and mesencephalic nuclei (yellow, ±200 ms windows, 5-70 Hz band, pseudo-t = 0.3), vii. deep mesencephalic n. (±200 ms windows, 5-70 Hz band, pseudo-t = 0.3), viii. substantia nigra (±200 ms windows, 5-70 Hz band, pseudo-t = 0.3), ix. tegmental n. (±200 ms windows, 5-70 Hz band, pseudo-t = 0.3), x. pontine or mesencephalic n. (±200 ms windows, 5-70 Hz band, pseudo-t = 0.3), and xi., and xii. all peaks in primary motor cortex (area 4, ±300 ms windows, 4-70 Hz band, all pseudo-t’s = 0.2).

Similarly, left hippocampal stimulation also elicited activity throughout the temporal network. Functional hippocampal and temporal networks engaged by a 20 Hz optical sine input to left hippocampus (as seen in Figures 3ai-iv) are shown for the third NHP in Figure 5a. Whole-brain SAM SPMs for this stimulus exhibited peaks in the following areas: (i.) left insular proisocortex/temporopolar proisocortex (±4 s windows, 10-30 Hz band, pseudo-t = 2.3), (ii.) left para-insular cortex (±4 s windows, 10-30 Hz band, pseudo-t = 2.4), (iii.) left dorsolateral amygdala (lateral n., ±4 s windows, 10-30 Hz band, pseudo-t = 2.0), (iv.) left basolateral/lateral amygdala (±4 s windows, 10-30 Hz band, pseudo-t = 2.0), (v.) left hippocampus, site of stimulation (±4 s windows, 10-30 Hz band, pseudo-t = 1.7), (vi.) left parasubiculum/presubiculum of hippocampus (±4 s windows, 10-30 Hz band, pseudo-t = 1.6), (vii.) contralateral (right) hippocampus (±4 s windows, 10-30 Hz band, pseudo-t = 2.6), and (viii.) left pontine reticular activating formation (±4 s windows, 10-30 Hz band, pseudo-t = 2.0). A second example of hippocampal network activity in a different NHP (as seen in Figure 3biii) in response to 8 Hz sine wave input is presented in Figure 5b. The SAM peaks elicited by this stimulus are as follows: (i.) left insula (±2 s windows, DC-15 Hz band, pseudo-t = 1.0), (ii.) PGa/IPa association areas of left temporal cortex (lateral yellow peak, ±2 s windows, DC-15 Hz band, pseudo-t = 1.0), left hippocampus (green peak, site of stimulation, ±2 s windows, DC-15 Hz band, pseudo-t = 1.0), and possible left red n. (yellow, medial peak, ±2 s windows, DC-15 Hz band, pseudo-t = 0.6), (iii.) area TEA/TEM of left temporal cortex (±2 s windows, DC-15 Hz band, pseudo-t = 1.0), and (iv.) area TL/TFM of right temporal cortex (±2 s windows, DC-15 Hz band, pseudo-t = 0.9). A third example of hippocampal network activity in response to 40 Hz sine wave input (same subject as in Figure 5b) is presented in Figure 5c. The SAM peaks are as follows: (i.) left anterior hippocampus (±1.5 s windows, 20-50 Hz band, pseudo-t = 0.9), (ii.) left hippocampus, site of stimulation (±2 s windows, 20-55 Hz band, pseudo-t = 0.6), (iii.) possible left deep mesencephalic n. (±2 s windows, 20-55 Hz band, pseudo-t = 0.7), and (iv.) left pulvinar n. of thalamus (±1.5 s windows, 20-50 Hz band, pseudo-t = 0.4). Collectively, these results indicate the utility of combining optogenetic techniques with MEG for functional brain mapping and mapping of deep structures, even under anesthesia.

## DISCUSSION

This study represents the first successful combination of optogenetics with magnetic source imaging of biomagnetic signals in non-human primates, and provides additional information on the relationship between MEG signals and the underlying electrical activity in the brain^28^. In this study, we have demonstrated anatomically-localized and temporally-precise control over neural activity in a MEG-compatible optogenetic preparation.

We provide evidence of the feasibility of optogenetic stimulation in the MEG environment and show that after optogenetic stimulation, both arcuate cortex and hippocampus can be localized by MSI across multiple NHPs, across stimulus types, and at a voxel size of 750 μm^3^. In addition, SAM can be used localize activity to NHP occipital cortex in response to visual stimulation. While the amplitude of the localized signal (as given by SAM pseudo-t score) varies, this is likely due to the type of stimulus input and can be used in the future to examine the effects of different time-varying inputs into a target structure. This study also provides evidence that a signal generated in deep brain structures is detectable with MEG, including hippocampus, and that downstream activation of other structures such as amygdala, caudate nucleus, putamen, globus pallidus, thalamic nuclei, superior colliculus, and even white matter is also detectable. Detection is possible even in the presence of high noise and suboptimal conditions resulting from the experimental preparation (e.g., the small size of the vervet brain, a large heartbeat artifact, artifact from anesthesia monitoring equipment, etc.).

Studies using non-human primates in MEG have previously demonstrated the feasibility of equivalent current dipole^32^ and beamformer^33^ analyses in monkeys. This study extends these observations to analysis of areas beyond superficial cortex into deep structures in an NHP species with a smaller brain, strongly supporting the validity of MSI in detecting deep brain signals. Furthermore, this study uses experimentally controlled, optogenetically generated signals from a known source and onset time to demonstrate the accuracy of localization. The current results suggest that a combination of whole head coverage with axial gradiometers, a low noise floor, low head movement, and appropriate use of beamformers allows deep localization of activity that is consistent with theoretical investigations and data from epilepsy patients^2–5,8,11^. SAM normalizes MEG signals with respect to the noise in the unique recording environment and with distance from the sensors; therefore, deep signals are localizable given sufficient signal-to-noise ratio (SNR). SNR amplification in this study was achieved through repeated presentations of the stimuli, by low head motion, and by beamforming itself, which also removes noise^7^.

The relevance of this finding to the clinical evaluation of epilepsy surgery candidates is significant. Currently, MEG is FDA approved for pre-surgical evaluation of epilepsy, but is often limited to dipole analyses only^16,34,35^. Unfortunately, dipole analyses break down with low SNR^5^, which is often the case with deep signals. The most common epilepsy type requiring surgical resection is mesial temporal lobe epilepsy wherein the seizure generator is deep in the brain^36^, thus methods permitting the localization of deep signals with MEG in such patients is of great utility in surgical planning. To this point, our results demonstrate that SAM can accurately localize a deep signal to its source (hippocampus). Furthermore, while Mikuni et. al. estimated 4 cm^2^ of active brain tissue is required for the localization of spikes^13^, our results show that significantly less tissue may be required to generate a detectable MEG signal, simply because the vervet hippocampus is far smaller, as are the structures comprising the CSPTC and visual networks. Additionally, the injection parameters of our viral vector^37-39^ and microscopy suggest that the volume activated was only a few mm^3^, and while it is likely that feed forward activation occurs we suspect that this falls well below 4 cm^2^ of tissue (see Figures 1ai, bi; Figure 3aiv, for examples).

The results of this study suggest that optogenetics can be combined with MEG imaging in non-human primates to permit precise functional mapping. Stimulation of arcuate sulcus activated the cortico-striato-pallido-thalamo-cortical (CSPTC) network such that voxels of peak activation fell within a functionally and anatomically described pathway^26,27^. Further, hippocampal stimulation evoked widespread activity in regions such as amygdala, insula, and contralateral hippocampus. Finally, natural visual stimulation using flashes of light elicited activity throughout the visual system. In the future, the combination of optogenetic techniques with MEG/MSI could be used to variably activate or inactivate discrete parts of the brain with excitatory or inhibitory opsins to reveal functional relationships that have not previously been described.

The temporal resolution of MEG enables this combination of techniques to map transient interactions that are not detectable using other functional imaging approaches^40^. Recent analysis techniques designed to increase fMRI sampling rate (e.g. dynamic functional magnetic resonance inverse imaging^41^), have achieved a maximum sampling rate around 10 Hz with a 5 mm^3^ voxel size^42,43^ with the 10 Hz period corresponding to events lasting 100 ms. In contrast, arcuate cortex was stimulated with trials of 50 ms square wave inputs and beamforming revealed arcuate peaks at 75 ms, and with a voxel size of 750 µm^3^ (Figure 2a), and 50 ms flashes of light elicited SAM peaks at 50 ms in occipital cortex (Figure S1aviii). These fast-transient events occur well beyond the temporal resolution of metabolically based methods and the recruitment of downstream areas can happen rapidly, suggesting that functional maps and connectivity analyses will vary depending on the duration and sampling rate of analyzed data. For example, fMRI network architecture has been demonstrated to change over short timescales (hours/minutes) during learning^44^, while analysis of MEG based network architecture has demonstrated variability on the order of milliseconds^45^. It is clear from this demonstration that the high sampling rates of MEG coupled with SAM affords exceptional advantages over traditional neuroimaging techniques. However because the beamformer problem is without a unique solution there will always be inherent uncertainties in localization^46^ without “ground truth” experiments utilizing controlled inputs to known brain targets. To this end, the approach we report may assist in efforts to continually to improve inverse problem solutions to yield greater certainty^30,47^.

In addition to the measurement of functional circuits, optogenetic control of neuronal field potentials combined with MEG will make it possible to investigate relationships that exist due to specific, transient activity patterns. For example, it may be possible to capture native activity at a recording site, use optogenetic stimulation to approximate population activity, and map the resultant response irrespective of state or behavior^48^. We anticipate that experimenter control over discrete brain regions of interest coupled to whole brain recording and detection of downstream responders will provide a platform for improved investigation of brain function in both normal and pathological conditions. Subthreshold modulation could be used to investigate population activity that gives rise to neural oscillations^49,50^ and suprathreshold synchronous modulation could be used to mimic epileptiform discharges in the study of epilepsy^50^.

The results of this study support, through direct measurement, the assumed theoretical ability of MEG/MSI to detect and localize experimentally controlled activity in the brain at very fine temporal and spatial resolution. We have demonstrated the feasibility of localizing activity in the brains of vervet monkeys using whole head instrumentation designed for human subjects. By combining optogenetic stimulation in MEG with beamformer analysis it is possible to empirically determine the practical abilities and limitations of MEG/MSI. Additionally, it is clear that this combination of techniques may permit functional brain mapping in spatial and temporal domains not otherwise possible with current functional imaging techniques.

## ACKNOWLEDGEMENTS

CC was supported by NIH grant R01 MH097695. JBD was supported by NIH grant NIAAA R21 AA028795, P01AA021099-S1 and UL1TR001420. DG was supported by NIH Grant NIAAA R01AA016852, P50AA026117, W81XWH-13-2-0095, NINDS 1R21NS116519, and UL1TR001420. DK was supported by NIH grants NIAAA R01AA016852, NINDS 1RO1NS105005, and NINDS 1R21NS116519. GA was supported by NIAAA F30 AA 23708-02 and NINDS NS073553. JSK was supported by VA/DoD W81XWH-13-2-0095. We wish to thank the Department of Neurology and Dr. Gautam Popli for providing scanner time on the MEG and the Translational Imaging Program (then Center for Biomolecular Imaging) for MRI pilot scans. We acknowledge the Wake Forest University Primate Center for providing animals for this study OD010965 (Jay Kaplan) and the Wake Forest Biology Microscopic Imaging Core Facility. We also acknowledge support from the W.G. (Bill) Hefner Veterans Affairs Medical Center and VA Mid-Atlantic Mental Illness, Research, Education, and Clinical Center. Finally, we would like to thank Dr. Caroline Bass for providing the viral construct used in this study.

## ONLINE METHODS

### Surgical Targeting

All studies were approved by the Animal Care and Use Committee of Wake Forest School of Medicine. This study was repeated in three female vervet monkeys, *Chlorocebus aethiops*, aged 15-16 years and weighing between 4.99-6.75 kg. Each animal was mounted in an MRI compatible stereotaxic frame with lipid containing ear bars, and T1-weighted anatomical images were acquired using a 3D-MPRAGE 0.5 mm isotropic sequence in a Siemens 3 T Skyra system with a custom built 8-channel flexible head coil (Dr. Cecil Hayes, Univ. Washington). The animals were removed from the stereotaxic frame and an additional structural MRI scan was acquired in which the animals were fitted with three lipid biomarkers fixed in the location of the MEG fiducials (placed in the conventional three-point fiducial locations) for co-registration with MEG data. The locations of the MEG fiducials were tattooed on the animals prior to the start of the study to ensure precise co-registration across multiple sessions. In one animal a post-mortem MRI was conducted after implantation and recording to confirm stereotaxic targeting (Figures. 1aii-iii and bii-iii).

Using Medical Image Processing, Analysis, and Visualization (MIPAV) software available from the NIH, stereotaxic coordinates were generated for CA3 of left hippocampus and for the posterior wall of the contralateral, right arcuate sulcus.

### Surgery

Animals were securely mounted and centered in a surgical stereotaxic frame. The cranium was exposed and ten ceramic screws (Rogue Research Inc.; Montreal, Canada) were placed around the perimeter of the surgical margin. A craniotomy (∼17 mm) was then drilled over the coordinates for each targeted brain region. A custom polyether ether ketone (PEEK) cannula with a Teflon stop was advanced through the dura on a stainless steel stylette. The stylette was retracted leaving the cannula in place; injection of the viral vector and placement of the optrode (optical fiber and recording electrode) was done through the cannula to ensure co-localization. A gas-tight, micro volume Hamilton Syringe was then filled with AAV10-CaMKIIa-ChR2-eYFP (provided by Dr. Caroline Bass, SUNY Buffalo) and 4 μL of virus was injected at each coordinate through a 38 G needle at a rate of 0.5 μL/minute. The syringe was left in place for 5 minutes following the injection and slowly removed. A custom made, chronically indwelling optrode was implanted to a depth of 0.5 mm above the injection coordinates. The optrode consisted of either a 200 or 400 µm diameter fiber optic cable (ThorLabs, Inc.; Newton, NJ) coupled to a 75 μm Pt/Ir electrode (FHC, Inc.; Bowdoin, ME) or a 35 μm formvar-coated Tungsten electrode (California Fine Wire Co.; Grover Beach, CA), which extended 0.7 mm beyond the tip of the optical fiber. Lastly, a custom (PEEK) headwell with a removable cap (Crist Instruments Co., Inc.; Bethesda, MD) was fitted over each craniotomy and cemented in place. The craniotomy was sealed with bone cement securing the optrode in place.

To ensure electrical and magnetic silence, we avoided the photovoltaic effect by designing optrodes with a beveled tip so that the conductive surface was in shadow rather than in the cone of light^22–24^. The absence of artifact was confirmed in saline and in pre-expression control recordings.

### Control for Photoelectric Effects

Prior to implantation, a combined fiberoptic/recording electrode (optrode) was submerged in saline and LED stimulation was conducted to rule-out artifacts associated with the photovoltaic effect. No transient artifact associated with the LED was detected upon stimulation and there was no significant difference in LFP amplitude from the pre-stimulation period suggesting the optrode design used was not susceptible to the photovoltaic effect.

### Electrophysiology Recordings

Recordings were conducted under anesthesia, using propofol (200-400 μg/kg/min) for the MEG scans and ketamine (12-15 mg/kg) and dexmedetomidine (0.0075-0.015 mg/kg) for the electrophysiological recordings. MEG recordings were conducted in a magnetically shielded room (MSR; Vacuumschmelze GmbH & Co.; Hanau, Germany) with a CTF MEG™ whole cortex helmet equipped with 275 first-order axial gradiometer coils, each with a 5 cm baseline and 22.4 mm average inter-sensor spacing and 29 reference sensors (CTF MEG International Services Limited Partnership; Coquitlam, BC, Canada). MEG recordings were sampled at 2400 Hz for a bandwidth of DC-600 Hz. Data were powerline filtered offline for 60 Hz harmonics with a 4 Hz width, and synthetic third order gradiometry was applied. Simultaneously recorded LFP data were amplified using a Nicolet intraoperative monitoring system (Natus Medical; Pleasanton, CA) sampled at 5 kHz with maximum range of +/- 5 mV with a bandpass filter of 0.1 Hz to 3 kHz and recorded in line with MEG data. On separate days without MEG recordings, LFP data were recorded using a SciWorks recording system (DataWave Technologies; Loveland, CO), AM-3600 extracellular amplifiers (A-M Systems; Carlsborg, WA), and a T8G100 headstage amplifier (Triad Biosystems International; Durham, NC) and bandpass filtered at 0.3 Hz to 5 kHz, with a gain of 500 and a 40 kHz sampling rate.

Prior to the optrode implantation, 8-minute resting-state MEG scans were acquired to establish baseline activity. Two weeks after surgery (before high level transgene expression, and after the animal had recovered from surgery) animals were stimulated for the first time at a range of optical intensities (10.3-75.6 mW/mm^2^ for 200 µm and 2.6-18.9 mW/mm^2^ for 400 µm fiber). These recordings were conducted in the MEG MSR under anesthesia as described, as well as in separate sessions in a dedicated electrophysiology suite. A variety of optical stimuli were delivered, including single 10 ms or 50 ms square pulses of 473 nm light from either a light emitting diode (ThorLabs) or from a laser (Shanghai Dream Laser Technology Co., Ltd; Song Jiang, Shanghai, China), 20 Hz square wave pulse trains, single triangular (sawtooth) ramps, and sine waves at 8, 20, or 40 Hz (see below for more stimulus details). Pulse duration of the laser was controlled with a 2 mm Uniblitz laser shutter and a D880C Uni-stable driver (Uniblitz Electronic; Rochester, NY). LED pulse durations were controlled with custom MATLAB scripts (Natick, Massachusetts: The MathWorks Inc.) and a digital-analog converter (DAC; Data Translation, Inc.; Marlborough, MA). After stable transgene expression was determined, the animals were stimulated again using the same paradigms as above and the resulting activity was recorded. Electrophysiological and MEG recordings after stable expression were again done simultaneously using the Nicolet system for acquiring electrophysiology during MEG and also separately.

The optrodes were tested in saline prior to implantation; *in vitro* two weeks after transfection; as well as post mortem. As a positive control for the electrode and MSI, analysis of visual stimulation was conducted. One hundred nineteen, 50 ms pulses of white light were presented via fiber optic cable to the left lower quadrant of the visual field of the left eye of the sedated animal. The right eye was taped shut per typical recording protocol while the left eye remained open. The fiber optic cable was channeled into the helmet slightly lateral and inferior to the left eye. The light pulses had a 3 s ISI via LED, and the experiment was performed in both the electrophysiology suite and in the MEG MSR. Timing was controlled by custom MATLAB scripts.

### Analysis

All analyzed data were derived from recordings in which motion was no greater than 0.2 mm. Head motion was nearly eliminated through the use of anesthesia and a carefully supported head.

Data preprocessing, head model creation, and beamforming were performed using CTF MEG™ Software (CTF MEG International Services Limited Partnership, Coquitlam, BC, Canada). MEG preprocessing included DC-offsetting, application of synthetic third order gradiometry, and powerline filtering^7^. MEG data were then co-registered with the monkey’s anatomical MRI data using the standard three-point fiducials. From this, a multiple-overlapping-spheres model of the head and whole brain volume was generated^25^. Dual-state SAM^3,5^ was applied to each active time segment and an equal length control segment composed of data preceding the stimulus, to construct 3D images of the entire brain volume. Noise normalized t-deviate statistical parametric maps of source power were derived from the beamformer output at a voxel size ranging from 0.75 - 1.5 mm^3^. Results are shown for 750 µm^3^ analyses due to superior accuracy in source localization. Such small voxel sizes were possible because of extremely low motion associated with this preparation. The degree of activation in each voxel is indicated by a pseudo t-score. Local maxima (synchronization) and minima (desynchronization) of the SAM maps were identified as voxels of peak activity.

SAM time-frequency analysis parameters were determined in part by the time-frequency characteristics of the input stimulus and by the power spectral densities (PSDs) and timing of the neural response measured on the LFP during simultaneous MEG/LFP recordings in the MSR when such recordings were obtained. In general, for brief stimuli, the beamforming windows necessary to detect activity changes also needed to be short, with a correspondingly wider frequency band to provide enough samples for analysis. For the sine wave inputs, the SAM frequency bands were set to bracket the sine frequency, and because the sine inputs were of longer duration the beamforming windows were correspondingly longer, both to provide enough samples for the analysis and also because the brain circuits appeared to sustain activity for a long time (see Results). Specific arcuate beamforming parameters for each stimulus type are as follows. For the 50 ms square pulses (120 trials, 5 s ISI), dual-state SAM relative to stimulus onset was conducted in 50 ms steps from 100-500 ms inclusive and for another window of 75 ms (Figures 2ai-iv). For each time window, the high-pass was set above the threshold for aliasing and the low-passes were varied in a series of 10 Hz steps ranging from 40-100 Hz and another step at 35 Hz. Peak voxels were identified in each whole-brain, time-frequency map and because peak voxels often appeared in multiple maps the time-frequency combination that expressed the maximal pseudo-t score for each peak was chosen as the representative map. A primate neuroanatomist (J.D.) confirmed the neuroanatomic localizations of the peaks to their target structures and also for downstream structures that were also activated following stimulation. For the 8 Hz sine waves (3 s stimulus length, 20 trials, 10 s ISI, Figure 2bi) the SAM time-frequency parameters were 1-4 s windows in 500 ms steps, a DC high-pass, and low-pass steps of 12, 15, 20, and 50 Hz. For the 40 Hz sine waves (3 s stimulus length, 20 trials, 10 s ISI, Figure 2bii) the SAM parameters were 500 ms - 4 s windows in 500 ms steps, a high-pass series ranging from 10-35 Hz in 5 Hz steps, and low-pass steps at 50 and 55 Hz. For the 10 ms square pulses (100 trials, 6 s ISI, Figure 2biii), SAM was conducted in 50 ms steps for windows from 100-750 ms, plus additional windows at 125 ms and 1 s, while the frequency bands had a minimum high-pass set to prevent aliasing for each time window, and a low-pass set at 50, 70, and 100 Hz. For the 20 Hz square pulse train (5 ms width squares, 10 pulses/train, 100 trains, 6 s ISI, Figure 2biv) the SAM time frequency parameters included windows from 100-500 ms conducted in 100 ms steps, plus an extra window at 750 ms, a high-pass set to prevent aliasing for each window, and a low-pass at either 70 or 100 Hz.

Beamforming parameters for the hippocampal stimulation experiments are as follows. For the single 60 s, 20 Hz sine wave (5 V, Figure 3ai-iv) the dual-state SAM time windows were 2, 3, 4, 5, 10, 20, 30, and 60 s, the high-pass was either 10 or 15 Hz, and the low-pass was 30 Hz. For the single sawtooth ramps (150 ms duration, 20 trials at an intensity of 4.996 V and 9.528 W, 6 s ISI, Figure 3bi) the SAM parameters included 150-400 ms time windows in 50 ms steps, plus additional windows at 125 ms and 500 ms, a high-pass set to prevent aliasing per time window, and low-passes ranging from 40-55 Hz in 5 Hz steps, plus additional windows at 70, 100, and 120 Hz, plus at 35 Hz for windows of sufficiently long duration. For the 10 ms square pulse stimuli (114 trials, 4.996 V/9.28 W, 6 s ISI, Figure 3bii) the SAM time-frequency parameters included 50, 175 ms windows, in 25 ms steps, a high-pass set to prevent aliasing per time window, and low-passes ranging from 30-100 Hz (when the time window permitted a small low pass) in 5 Hz steps, with additional windows at 120 and 200 Hz. For the 8 Hz sine waves (3 s duration, 20 trials, 10 s ISI, Figure 3biii) the beamforming parameters included 1-3 s windows in 500 ms steps, a DC high-pass, and a low-pass of 12, 15, or 20 Hz. For the 40 Hz sine waves (3 s duration, 20 trials, 10 s ISI, Figure 3biv) the beamforming parameters included 1-3 s windows in 500 ms steps, high-passes of 10, 20, 30 or 35 Hz, and low-passes of 50, 55, or 60 Hz.

Beamforming for the visual stimulation experiment (Figure S1) included 50-300 ms windows in 25 ms steps, a high-pass set to prevent aliasing per time window, and low-passes ranging from 30-70 Hz in 5 Hz steps, plus additional windows at 80, 90, 100, 120, and 200 Hz.

LFP data were analyzed using SciWorks, Neuroexplorer (Nex Technologies, Madison, Al), and custom scripts in MATLAB. Peak amplitude of the LFP in response to optical stimulation during electrophysiology recordings was measured between two-week, five-week, and seven-week time points.Peak amplitudes for each replicate were measured at the time point corresponding to the peak of the average trace for each light intensity.

### Histology

To confirm the anatomical position of the optrode placement and injection site, at the completion of the imaging study, one animal was necropsied by first sedating with ketamine followed by an overdose with pentobarbital. A thoracotomy was conducted and the animal was transcardially perfused with ice-cold phosphate buffered saline followed by 4% paraformaldehyde fixative. The brain was extracted and sectioned at a 50 µm thickness on a modified Vibratome 1000 (Leica Biosystems Inc.; Buffalo Grove, IL). Sections of tissue were mounted on gelatin coated microscope slides and imaged on a Zeiss LSM 710 confocal microscope with 10X and 20X objectives (Carl Zeiss Microscopy, LLC; Thornwood, NY).

---

**Supplementary Figure 1.**
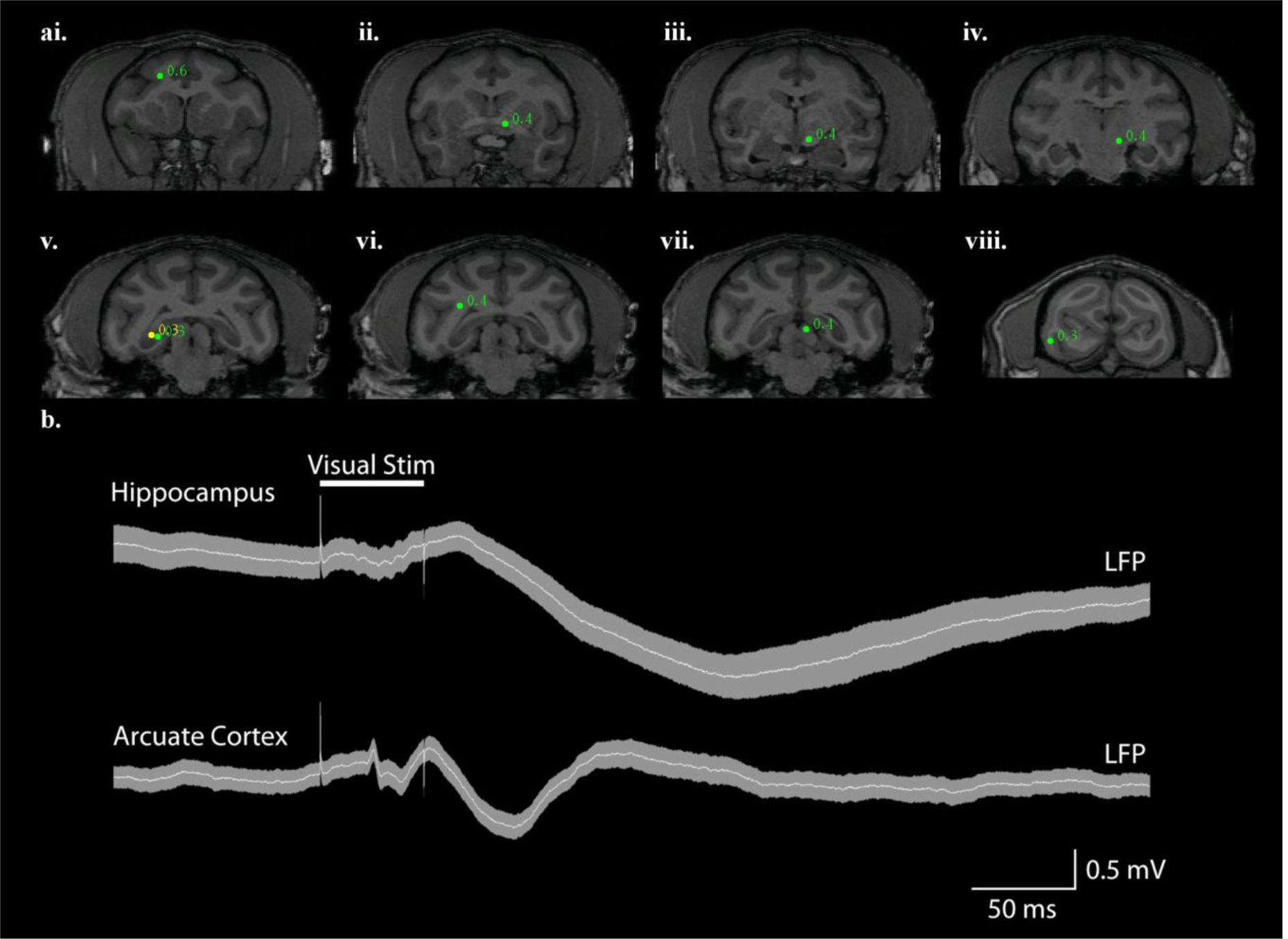
Visual stimulation engages related brain regions. **a**. Identification of functional networks in response to 50 ms light presentations to the left lower visual quadrant of the left eye. SAM analysis of the visual stimulus reveals both peaks in the known visual network as well as additional peaks, including: **i**. right anterior arcuate sulcus, **ii**. left anterior commissure, **iii**. left optic tract, **iv**. left lateral geniculate n. **v**. right posterior hippocampus, **vi**. right posterior white matter tracts, **vii**. left superior colliculus, and **viii**. right occipital cortex. **b**. LFPs recorded from right arcuate cortex and left hippocampus revealed clear event-related potentials in response to visual stimulation.

